# Comparative 3D analysis reveals species-specific patterns of coral polyp morphology and gastrovascular integration

**DOI:** 10.64898/2026.07.10.737875

**Authors:** Emma Rangel-Huerta, Meiru Wang, Stephanie H. Nowotarski, Keith Duncan, Sean McKinney, Matthew C. Gibson

## Abstract

Coral reefs are constructed by colonial cnidarians whose survival depends on the coordinated growth and physiological integration of thousands of interconnected polyps. While coral skeletons have been extensively studied, the internal three-dimensional organization of coral tissues remains poorly resolved, limiting our understanding of how reef-building corals function as integrated modular organisms. In this study, we established a contrast-enhanced X-ray tomography (XRT) workflow for decalcified coral tissues, enabling detailed visualization and quantitative comparison of internal polyp architecture across four reef-building species with distinct colony forms: *Acropora cervicornis, Acropora millepora, Montipora capitata,* and *Pocillopora damicornis*. Importantly, this methodology resolved previously inaccessible patterns of tissue organization and structural connectivity among neighboring polyps. The two *Acropora* species shared a conserved axial – radial organization but differed in mesenterial morphology, whereas *M. capitata* exhibited complex, entangled mesenterial networks that connected both neighboring and distant polyps. In contrast, *P. damicornis* displayed superficial connectivity restricted to the coenosarc. Together, these results suggest that internal tissue architecture is an evolutionarily flexible trait, shaped by ecological and developmental pressures rather than strictly by shared ancestry. Our XRT workflow thus provides a new comparative framework for understanding how corals function as integrated living colonies.

## Introduction

The ecological success of reef-building scleractinian corals depends on the integration of a multitude of genetically identical polyps acting as a single, integrated organism. Within coral colonies, each polyp is a biological unit comprising a mouth, tentacles, a pharynx, and a gastrovascular cavity enclosed within a sac-like body. Every polyp deposits calcium carbonate around itself, and the cumulative effect of this secretion across the colony is the formation of massive three-dimensional (3D) reef structures that support some of the most biodiverse marine ecosystems on earth ^1^. Understanding how corals integrate polyp-level processes with colony-level architecture is therefore central to coral biology, and especially urgent as reefs face accelerating climate-driven declines ^2^.

The growth of coral colonies is particularly complex, as they expand through the continuous clonal budding of new polyps, while sexual reproduction occurs seasonally through broadcast spawning at the colony level ^3, 4^. This iterative developmental process generates colonies comprised of modular units whose collective organization gives rise to traits such as branching patterns, skeletal architecture, and internal polyp-polyp connectivity ^4, 5^. The resulting diversity of colony forms is therefore related to the interplay between intrinsic developmental programs ^5, 6^ and the ecological niche, physiological function, and hydrodynamic performance ^7, 8^.

Despite the critical importance of polyp organization and connectivity, our understanding of coral colony morphogenesis is largely derived from observations of the mineralized skeleton. For example, X-ray tomography (XRT) and related imaging approaches have enabled detailed quantification of corallite geometry, skeletal density, and growth banding across species ^9–16^. These studies have shown remarkable skeletal diversity but have largely left the role of the living soft tissue unresolved. Current live micro-CT studies have shown key aspects of coral growth and the dynamic processes underlying biomineralization in living colonies. Emphasizing the importance of polyp morphogenesis as a process that precedes and guides skeletal morphogenesis, highlighting the central role of living tissues in shaping coral skeletal architecture ^17^. However, despite these advances, most studies have focused on surface dynamics and skeletal development, leaving the 3D polyp morphology within the colony largely unexplored. In most cases, the dense aragonite exoskeleton strongly absorbs X-rays, masking signal from the soft tissue ^18, 19^. When the skeleton is removed through decalcification, the thin and fragile coenosarc (a continuous tissue layer connecting polyps across the colony surface) loses its structural support and often collapses, degrading native morphology before imaging can take place. Consequently, the internal polyp anatomy of scleractinian coral colonies has traditionally been examined through two-dimensional histology ^20, 21^, which inherently fails capture the (3D) connectivity that defines the colony as an integrated system.

Connectivity has been studied in other cnidarians, including octocorals and hydroids, where specialized junctional structures regulate gastrovascular flow and coordinate colony-wide physiology ^22, 23^. In contrast, our understanding of similar processes in stony coral colonies remains limited. Previous studies have shown that surface currents generated by cilia act in combination with mucus-mediated flows to enable polyp-to-polyp exchanges across the colony surface ^24, 25^. Internally, gastrovascular circulation distributes nutrients and metabolites derived from symbiont photosynthesis throughout the coenosarc ^26–28^.

Due to the challenges of imaging within an intricate and complex skeletal architecture, fundamental questions about coral colony integration remain poorly understood, particularly how polyps are spatially organized and how they communicate with each other. Furthermore, it remains unclear how internal colony architecture varies across species with contrasting growth forms and ecological strategies. To address these questions, novel methods are required to preserve and resolve soft tissue in 3D within intact coral fragments. Contrast-enhanced XRT, using treatments with Lugol’s iodine or phosphotungstic acid, has proven to be a powerful method for visualizing soft tissues in a wide range of organisms ^29–32^ and has recently been applied to octocorals ^22^. However, applying contrast-enhanced XRT to stony corals requires additional technical steps, including decalcification, stain penetration and sample embedding. As a result, many coral XRT workflows remove the soft tissue and focus on dry skeletal samples ^7, 8, 13, 19^. This provides excellent resolution of skeletal traits but leaves the associated soft tissue architecture unresolved.

In this work, we designed and optimized a contrast-enhanced XRT workflow for decalcified scleractinian tissues, applying it to four species with diverse colony morphologies: *Acropora cervicornis*, *Acropora millepora*, *Pocillopora damicornis*, and *Montipora capitata*. This methodology establishes a reliable pipeline for visualizing polyp architecture, tissue organization, and gastrovascular connectivity. Our approach further enables comparative analysis across species with divergent growth strategies and interprets the resulting structural patterns within an ecological, evolutionary, and developmental perspective that links polyp-level morphology to colony-level form and function. By bridging skeletal morphology and living tissue architecture, this work opens a new window onto the structural principles that underpin coral diversity and integration.

## Results

### An XRT-based workflow for resolving internal polyp architecture in decalcified coral tissues

We first sought to establish an XRT-based workflow to resolve soft tissue architecture in decalcified coral samples, with emphasis on optimizing sample preparation. The overall workflow is summarized in Figure 1A. Following sample collection, we identified and optimized critical preparation steps (highlighted with an asterisk in Figure 1A), including decalcification, contrast agent staining, and incubation time. These steps required extensive testing to achieve optimal tissue preservation, contrast enhancement, and imaging resolution.

**Figure 1.**
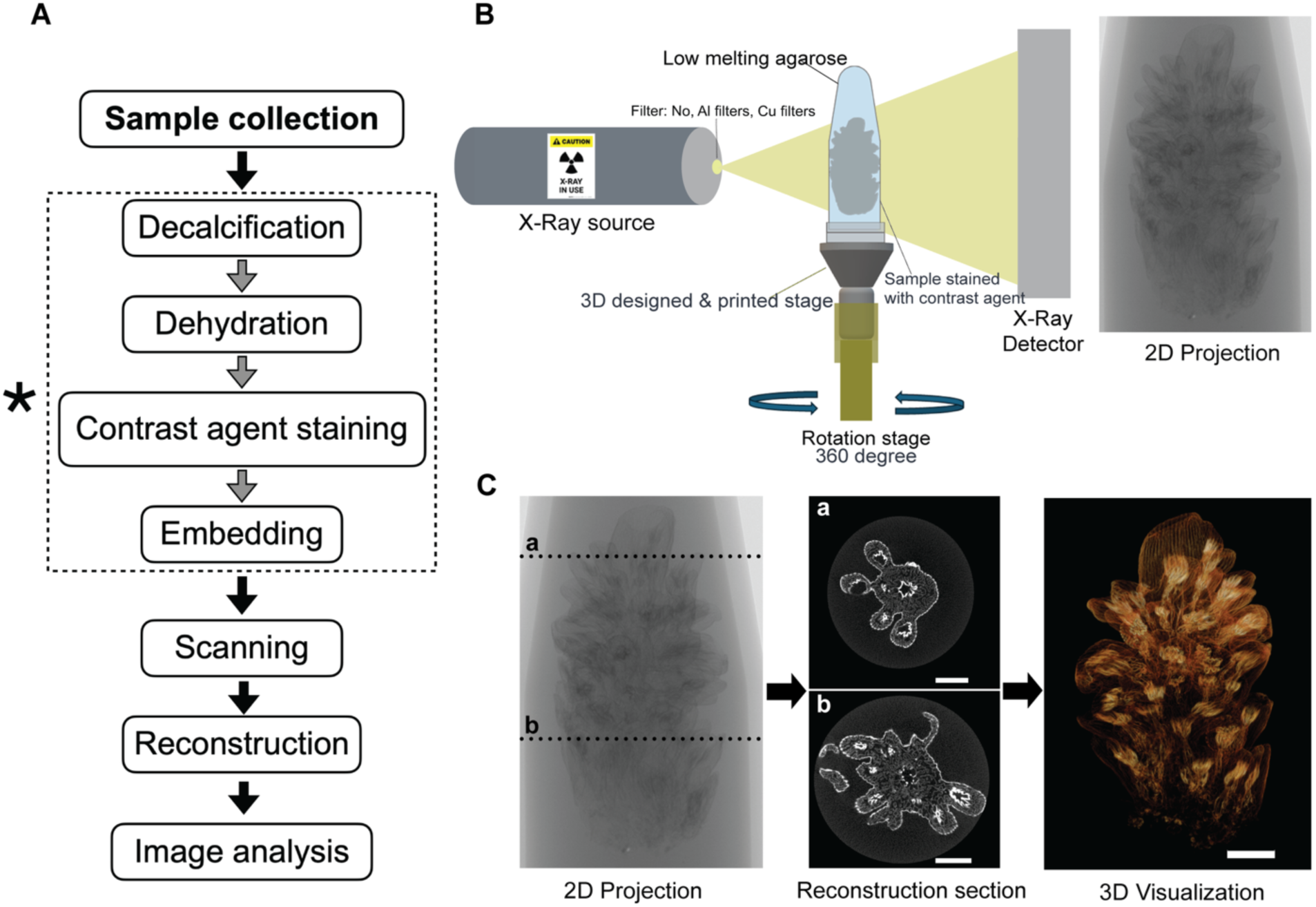
Contrast-enhanced X-ray tomography (XRT) workflow for coral soft-tissue imaging. **A,** Overview of the sample preparation and contrast-enhanced XRT workflow. After sample collection, key preparation steps, including decalcification, contrast-agent staining, and incubation time, were systematically optimized to improve tissue preservation, contrast enhancement, and imaging resolution. Critical optimized steps are indicated by asterisks. **B,** Schematic of the XRT scanning setup. The stained sample was immobilized in 1% low-melting agarose within a 1.5ml microcentrifuge tube. This tube was mounted onto on a custom 3D-printed holder and rotated 360° during X-ray acquisition to generate 2-dimensional projections. **C,** A pipeline from 2D projections to 3D soft-tissue visualization. Scale bars, 2000 μm.

The duration of decalcification varied among species, largely because skeletal density and fragment size varied considerably (Supplementary Table 1). We determined that decalcification was complete by a combination of visual inspection and mechanical verification. In most cases the samples were considered fully decalcified when they appeared evenly transparent under the dissecting microscope. Nevertheless, for species like *A. millepora*, which remained somewhat opaque even after mineral loss, we used a fine needle to puncture the base. If we detected no resistance, we considered them decalcified and used an identical timing for the rest of the batch. For corals that were attached to an aragonite substrate, such as *M. capitata*, the completion of decalcification was based on both transparency and tissue detachment from the substrate. After decalcification, what remained was a soft, porous tissue indicating that samples were ready to proceed with contrast-agent staining. Staining optimization was performed to maximize visual contrast for high quality 3D imaging. Following preparation, samples were embedded, mounted, and scanned using a micro-X-ray source (Figure 1B). During acquisition, each sample was rotated 0.2° per step for a total of 360°, yielding 1800 2-D projections per sample. The resulting projection series was then computationally reconstructed and rendered into 3D volumetric models, enabling visualization of internal polyp architecture (Figure 1C).

### Comparative colony morphology across four coral species

We first analyzed comparative 3D reconstructions of colony fragments from *Acropora cervicornis, Acropora millepora, Pocillopora damicornis,* and *Montipora capitata* colonies. This workflow revealed striking interspecific differences in polyp morphology, spatial organization, and polyp relationship with colony surface architecture (Figure 2). Both *A. cervicornis* and *A. millepora* colonies exhibited cylindrical branching morphologies; however, their internal organization differed markedly. In *A. cervicornis,* the branch tip was dominated by a large axial central corallite (*yellow arrow*), surrounded by elongated tubular radial corallites with outward-projecting margins (*white arrowheads*; Figure 2A). Reflecting this skeletal pattern, polyps were arranged around the dominant axial region and projected outward, producing an open and heterogeneous surface topology (Figure 2E, I). This configuration revealed a hierarchical organization with clear differentiation between axial and radial polyps (Supplementary Movie 1).

**Figure 2.**
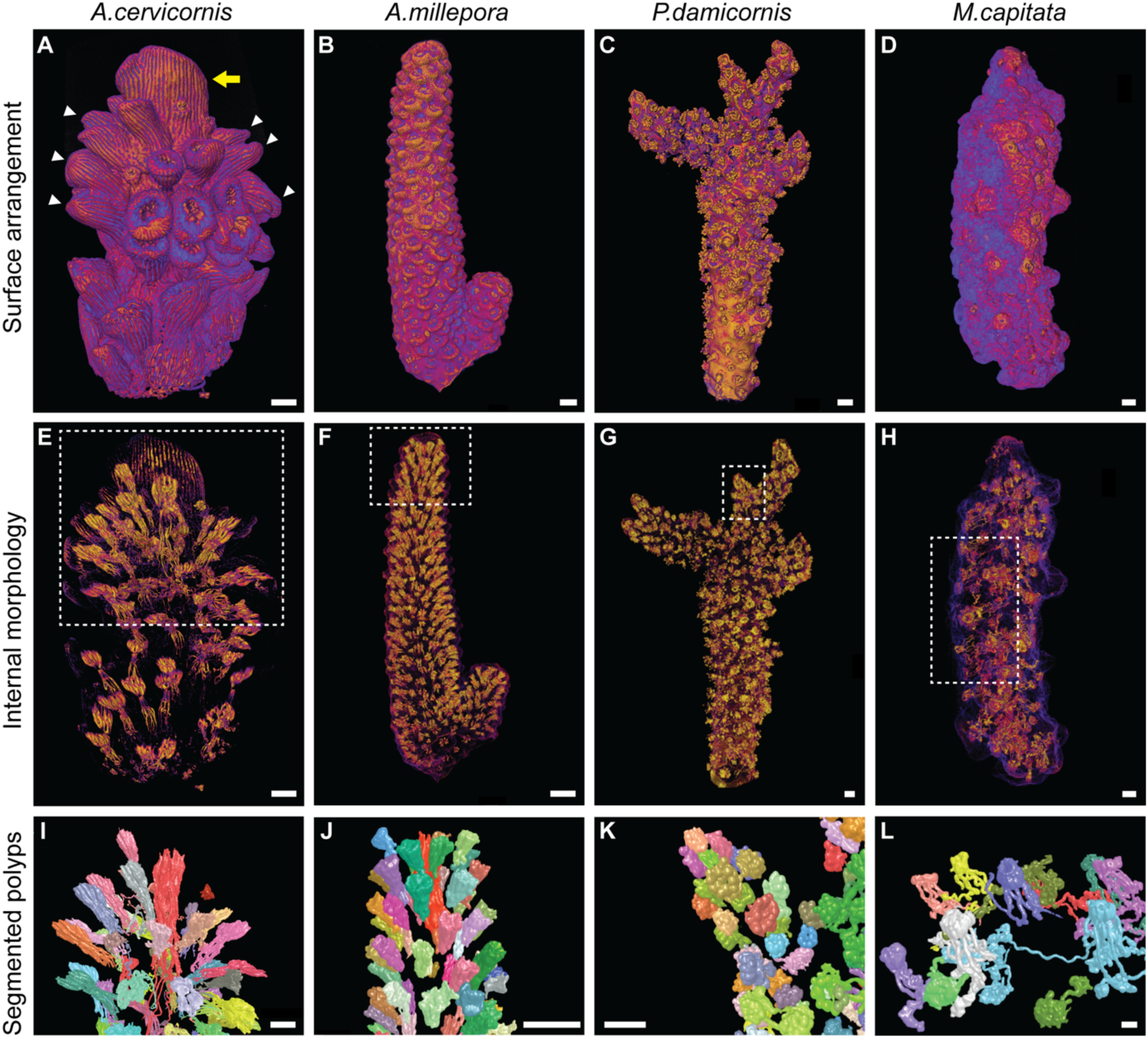
Species-specific differences in coral polyp arrangement and internal colony architecture. **A–D**, Volume renderings showing the overall surface arrangement of polyps in *Acropora cervicornis* (A); apical axis/axial region at the upper branch tip (yellow arrow) and surrounding lateral corallites (white arrowheads), *A. millepora* (B), *Pocillopora damicornis* (C), and *Montipora capitata* (D). **E–H**, Volume renderings highlighting internal morphology and spatial organization of polyps within the colony. Dashed boxes mark representative regions selected for segmentation. **I–L,** Representative segmented polyps from the boxed regions in E–H, showing variation in polyp morphology, and patterns of association among neighboring polyps across species. Scale bars, 1000μm.

In contrast, *A. millepora* exhibited a less prominent central axial corallite and radial corallites were shorter, more densely packed, and closely appressed to the branch rather than projecting outward (Figure 2B). These polyps were organized in a consistent outward-facing orientation, resulting in a more regular and homogeneous surface pattern (Figure 2F, J). This compact organization suggests a more uniform distribution of growth and tissue integration than that observed in *A. cervicornis* (Supplementary Movie 2).

*P. damicornis* colonies presented a highly branched, bush-like morphology with a dense and irregular 3D-shape. Corallites were small, circular, and packed tightly across the colony surface without any obvious spatial pattern (Figure 2C). Individual polyps were also relatively small (mean total polyp length = 1.17 mm), characterized by bearing twelve stump-like tentacles and shorter mesenteries compared to the Acropora species, with no elongated projections extending into the lumen. Rather than being recessed, polyps protruded from the surface, giving the colony a highly textured, rugose appearance (Figure 2G, K) (Supplementary Movie 3).

*M. capitata*, despite belonging to the same family as the two *Acropora* species (Acroporidae), exhibited notably distinct colony structure ^3, 33^. Its corallites were not raised cups, but low-relief, ring-like structures embedded within the coenosteum (Figure 2D). The colony surface was bumpy and irregular, with pronounced verrucae formed by coenosteum tuberculae (Figure 2D). Polyps were recessed within this structure with only the oral disk visible at the surface, and almost no protrusion (Figure 2D, H). Although no corallites were present in *M. capitata,* polyps showed surprising internal complexity, with mesenteries forming elaborate interconnected networks that extended throughout the fragment. Differences in polyp organization across species reflect distinct developmental patterns, whereby local tissue organization scales to shape the overall architecture of the colony.

Although *A. cervicornis* and *A. millepora* shared a conserved axial–radial organization, characterized by a dominant axial polyp surrounded by secondary radial polyps (Figure 3A, B, E, F), their internal organization differed markedly. We turned to digital segmentation, which involves manually tracing and isolating individual structures from the reconstructed 3D volumes. In *A. cervicornis*, the axial polyp displayed pronounced variation in mesenterial length, with some mesenteries extending along the main axis up to approximately 8.9 mm, whereas radial polyps exhibited shorter mesenteries of approximately 2.5 mm (Figure 3C, D). In contrast, *A. millepora* had smaller axial polyps (approximately 2.2 mm), with uniformly short mesenteries that did not extend along the axial lumen, while its radial polyps (approximately 1.3 mm) were also shorter than those of *A. cervicornis* (Figure 3G, H).

**Figure 3.**
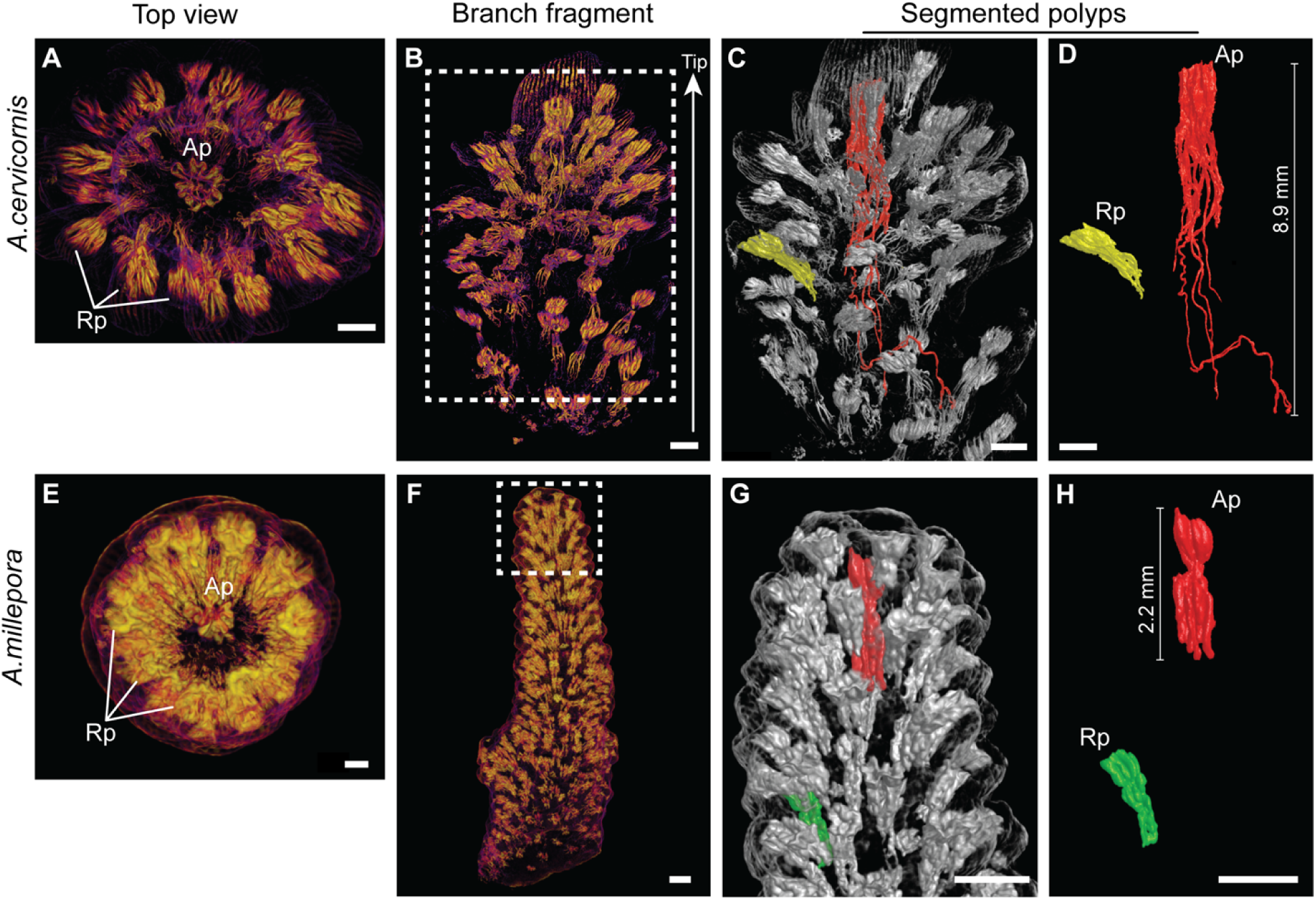
Contrasting branch-tip morphologies between *A. cervicornis* and *A. millepora.* Branch fragments of *A. cervicornis* (**A–D**) and *A. millepora* (**E–H**). **A, E**, Top-view volume renderings of branch tips showing the Axial polyp (Ap) at the apex surrounded by Radial polyps (Rp). **B, F,** Longitudinal volume renderings; dashed boxes indicate regions selected for segmentation. **C, G**, Representative segmented polyps shown within semi-transparent branch volume rendering; the axial polyp is shown in red and radial polyps in yellow (*A. cervicornis*) or green (*A. millepora*). **D, H,** Isolated segmented polyps highlighting the greater elongation of the axial polyp relative to radial polyps. Scale bars, 1000 μm.

In contrast to *Acropora* species, *M. capitata* does not exhibit an axis-driven growth pattern and notably lacked a clearly defined dominant axial polyp. Instead, polyps displayed substantial variability in total length and mesentery development. Some polyps exhibited pronounced projections extending deeper into the tissue, while others had shorter mesenteries with fewer and less extensive projections (Supplementary Movie 4). Together, these observations reveal fundamental differences in polyp hierarchy and internal organization across species, indicating that variation in developmental patterning underlies differences in colony growth strategies, integration, and functional organization.

Through our image analysis, we also identified early-stage polyps in *A. cervicornis*, allowing us to characterize morphological features associated with early polyp development (Figure 4). In this case, two newly emerging polyps were observed adjacent to the axial corallite (Figure 4A, *arrows*), characterized by a reduced calice compared to neighboring radial corallites. Internal views revealed a central opening corresponding to the mouth and six visible developing mesenteries extending from the body wall, with no clearly defined pharynx (Figure 4B, D). The internal lumen view exhibited elongated mesenteries extending along the axial lumen, whereas developing polyps showed shorter mesenteries and a simplified internal organization (Figure 4C; Supplementary Movie 5). These observations are consistent with a progressive model of polyp development, in which structural complexity increases as polyps mature and integrate into the colony.

**Figure 4.**
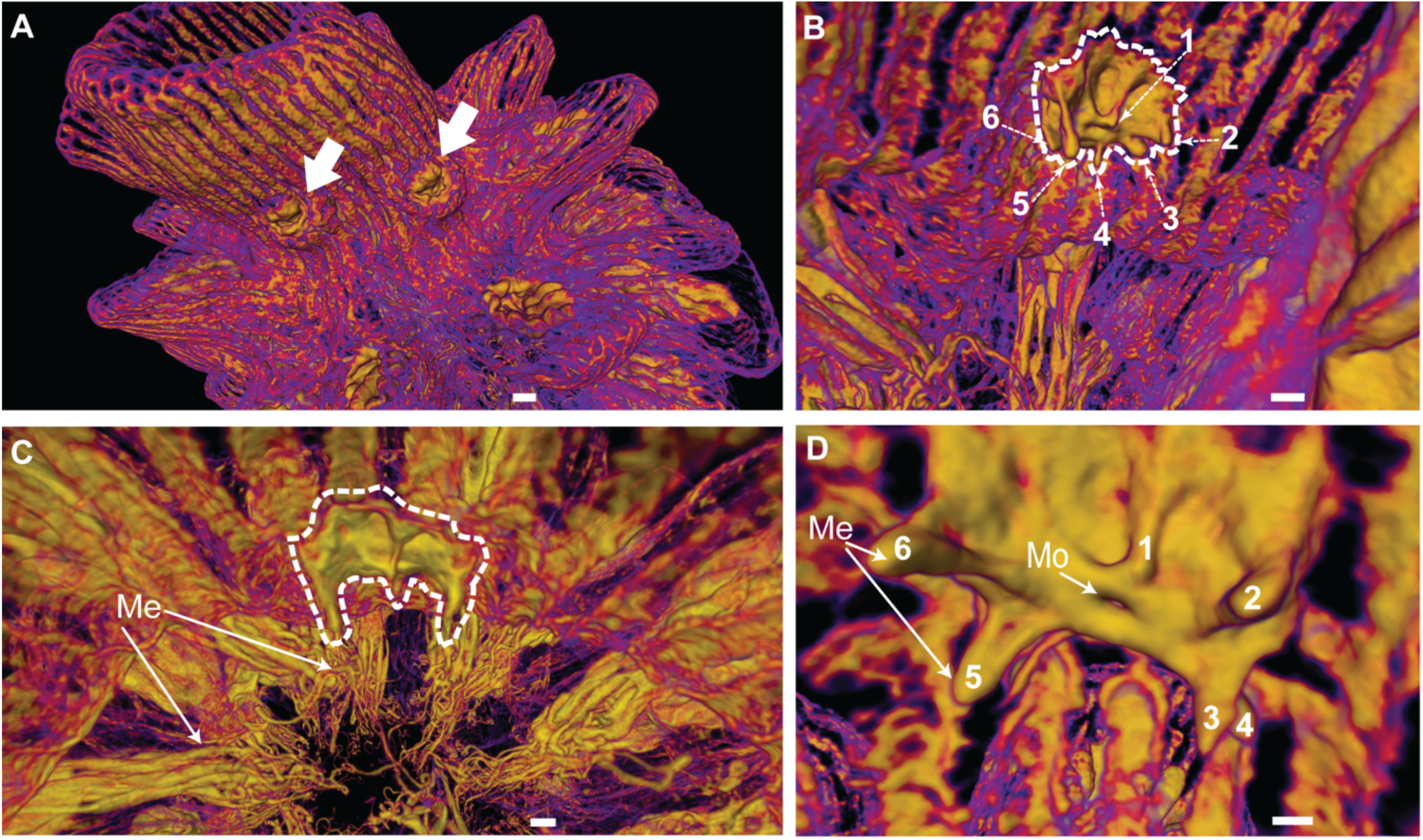
Coenosarc surface connects two newly emerging polyps at the branch tip of *A. cervicornis*. **A**, Longitudinal volume rendering of the apical region showing two newly emerging polyps (*arrows*) positioned laterally at the branch tip. Surface rendering showed both polyps with oral openings. **B, D,** High magnification of internal view of a newly emerging polyp showing six visible mesenteries (*Me*) and the oral opening (*Mo*). **C,** Internal view through the lumen showing mesenterial length in newly emerging polyps (*dashed lines*) and adjacent mature polyps, with neighboring mature polyps extending their mesenteries toward the main lumen. Scale bars, 1000 μm.

### Linking internal polyp organization to coral functional ecology and evolutionary diversification

Polyp morphology provides a fundamental framework for understanding coral colony development, as it reflects the processes by which polyps were generated, positioned, and integrated within a growing colony. Structural features such as mesentery length, orientation, and complexity offer key insights into tissue connectivity, resources distribution and physiological coordination. To assess interspecific variation, we next performed 3D reconstructions of individual polyps across species (Figure 5). We first defined the general polyp anatomy as a cylindrical, sac-like body with an upper central mouth surrounded by tentacles, a pharynx, and a gastrovascular cavity subdivided by vertical mesenteries, typically exhibiting hexaradial symmetry (Figure 5A).

**Figure 5.**
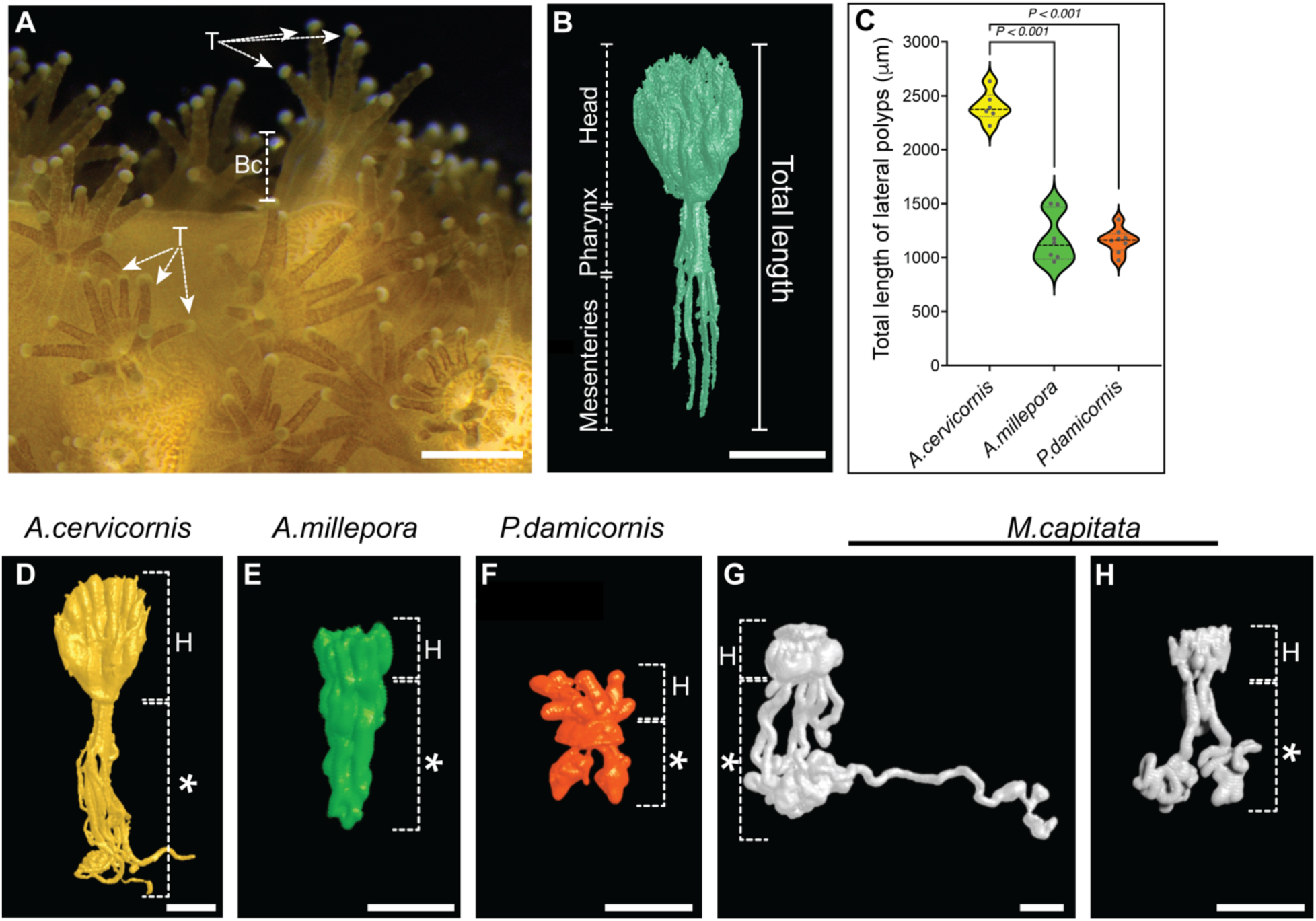
Divergent polyp architecture and mesenterial length across four coral species. **A**, Live colony view of. *P. damicornis* showing overall body plan of polyps across the colony; a body column (*Bc*) and extended tentacles (*T*) are indicated. **B,** Segmented polyp of *Acropora cervicornis* used as a reference to illustrate polyp anatomy and the total length measurement applied for comparative analysis across the four coral species. **C,** Violin plot showing polyp total length measured from the same individual. Data are presented as mean ± SD. Mean polyp total length was 2423 μm ± 139.8 (*n*= 6) in *A. cervicornis*, 1186 μm ± 224.8 (*n*= 7) in *A. millepora*; 1179 μm ± 114.1 (*n*= 8) in *P. damicornis*. Statistical analysis was performed using one-way ANOVA in GraphPad Prism. Statistically significant **** = *P* < 0.0001. **D–H,** Representative segmented polyps from the four coral species illustrating comparative differences in polyp morphology. Head (H) and pharynx and mesenteries (∗). Scale bars, 1000 μm.

Tentacle contraction is a well-known consequence of chemical fixation in corals. Despite the application of a MgCl_2_-based relaxation protocol, we found that tentacles were typically contracted within the region corresponding to the polyp head. However, since all samples underwent a similar fixation and decalcification protocol, tissue contraction and associated measurement limitations apply equally across all four species. A segmented polyp from *A. cervicornis* was used to define the anatomical regions for quantifying total polyp length (TL), including the oral pole (often regarded as the “head”, including the mouth, tentacles and pharynx), and mesenteries (Figure 5B). Comparative analyses revealed marked interspecific differences in polyp morphology. *A. cervicornis* exhibited elongated polyps with well-defined heads, a distinct pharynx, and extended mesenterial projections running parallel to the branch axis, reaching a mean total length of 2.42 mm (Figure 5C, D). In contrast, *A. millepora* displayed shorter, more compact polyps with reduced mesenterial extension and a more uniform internal organization (mean TL = 1.19 mm; Figure 5C, E). *P. damicornis* exhibited small, compact polyps with the shortest mesenterial projections of the four species, tapering to narrow, tooth-like tips rather than the broad, elongated extensions seen in Acropora (mean TL = 1.17mm; Figure 5C, F). Lastly, *M. capitata* showed pronounced variability in polyp morphology that precluded consistent measurement of total length across individuals. We identified two main structural patterns: centrally located polyps that exhibited densely entangled mesenteries with elaborate, interlaced projections, and peripheral polyps that displayed shorter mesenteries with less complex organization (Figure 5G, H; Supplementary Movie 6). These reconstructions reveal previously unresolved diversity in internal polyp architecture.

Importantly, our micro-CT workflow also enabled reconstruction of the gastrovascular architecture within coral fragments, revealing strikingly structural connectivity between polyps among *A. millepora*, *M. capitata* and *P. damicornis* (Figure 6). In *A. millepora*, volume rendering of the whole branch fragment provided an overview of the distribution of contrast-enhanced tissues and showed the crowded arrangement of polyps within the colony (Figure 6A). From this overview, internal views revealed and apparent network of interconnected pathways extending through the main lumen and linking neighboring polyps (Figure 6A, dashed lines). However, because this network was dense and difficult to segment, we acquired a higher-resolution X-ray tomography scan to examine these connections at higher detail. This allowed us to resolve and segment continuous tissue connections between the axial and radial polyps, confirming the present of direct structural links between polyps (Figure 6B & Supplementary Movie 7).

**Figure 6.**
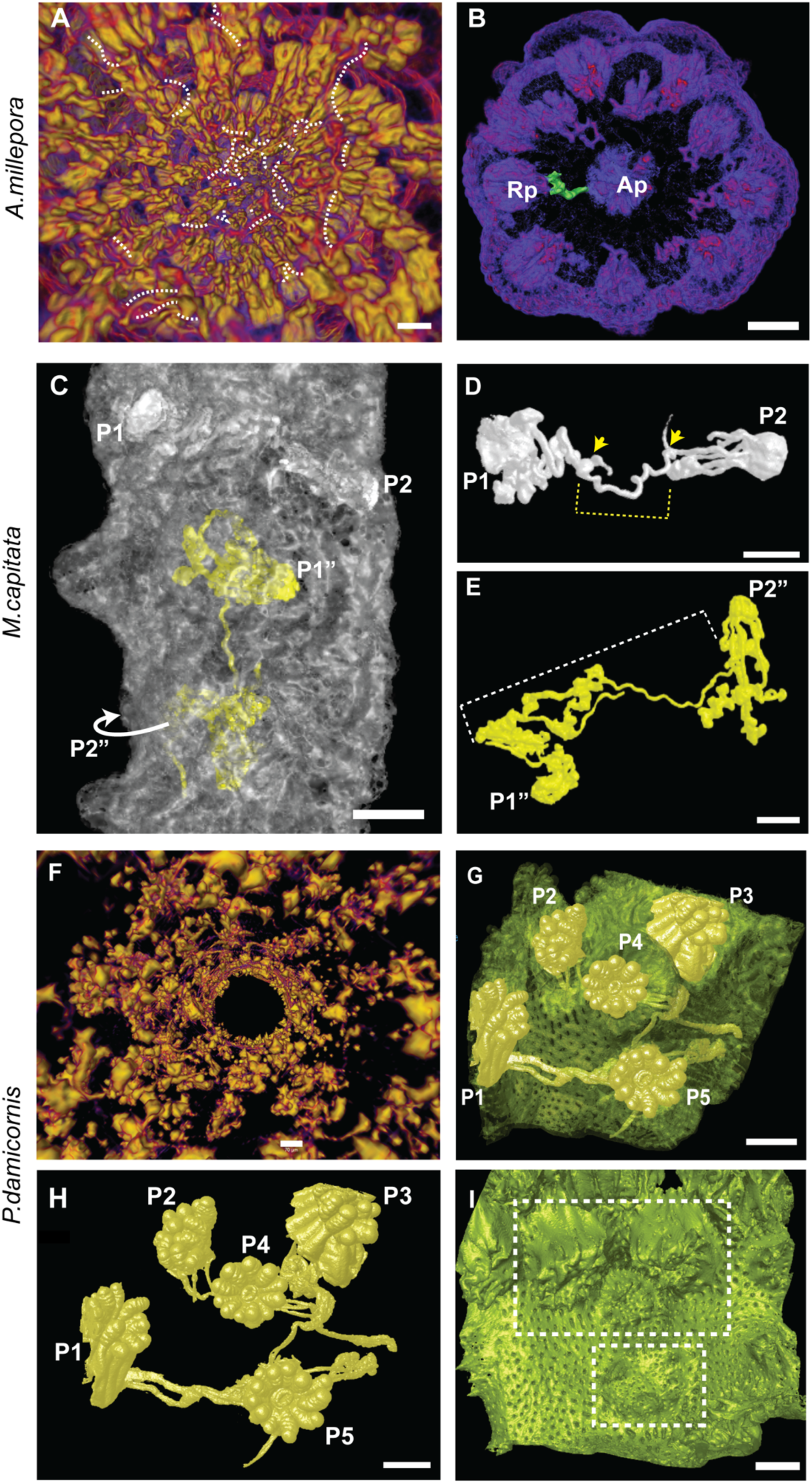
Coral architectures define divergent levels of internal polyp connectivity. **A, B**. *A. cervicornis* intra-colony network connections**. A,** Internal view of the mid-branch region, with neighboring polyps highlighted in yellow and putative connectivity pathways indicated by dashed lines. **B,** Cross-sectional view of the branch tip showing a structural link (*green*) between the Axial polyp (*Ap*) and Radial polyp (*Rp*), high magnification scan 4X. **C–E**. *M. capitata* segmented reconstruction resolving continuous structural connections between polyps and their relative position within the fragment. **C,** Transparent renderings of the *M. capitata* fragment, showing the spatial arrangement of nearby polyps P1-P2 (*white*) and non-adjacent polyps (*yellow*) P1”-P2” within the coral tissue (*gray*). **D,** Segmented reconstruction resolving a continuous connection between neighboring polyps through a mesenterial filament (*yellow dashed lines*); yellow arrows indicate the connecting pathway. **E,** Segmented reconstruction of two distant polyps, (connected through an intertwined mesenterial filament. **F–I**, *P. damicornis* displaying superficial polyp connectivity. **F**, Transparent volume rendering of *a* cross-section through the main axis of *P. damicornis*, showing a superficial pattern of polyp connectivity, with polyps highlighted in yellow. **G,** Segmented reconstruction of a polyp group, with polyps (*yellow*) and surface-level connections visible above the outer tissue layer (*green*). **H**, Isolated 3D reconstruction of segmented polyps and their connecting structures, highlighting links among polyps P1**–**P5. **I**, View of inter-polyp connectivity located beneath the coenosarc. Mesenteries display a dentition-like arrangement (*dashed rectangle lines*) oriented toward the main lumen, leaving an open tunnel along the main axis. Scale bars, 1000 μm except in **B**, 500 μm **F**, 70 μm.

Similarly, in *M. capitata*, volume rendering showed neighboring and distal polyps connected through internal tissue contacts (Figure 6C). We performed segmentation to further resolve these networks, identifying continuous structural links, including direct connections between their mesenteries, consistent with gastrovascular connectivity (Figure 6D). Surprisingly, these connections were not limited to immediate neighbors; individual polyps also formed links with more distant polyps across the coral fragment. This distance suggests that connectivity does not depend solely on spatial proximity (Figure 6E) & (Supplementary Movie 8).

*P. damicornis* is an imperforate coral, meaning its skeleton lacks the porous connections between corallites found in perforate species (Figure 6F). Resolving its connectivity patterns required higher resolution X-ray microscopy (XRM). Reconstruction of colony architecture showed more superficial polyp-to-polyp connections, just below the coenosarc, rather than an intricate network through the lumen (Figure 6G, H). We observed that the mesenteries themselves were short and terminated in pointed, tooth-like projections oriented toward the central gastrovascular cavity. These mesenteries did not extend into elongated structures like those observed in *Acropora* or *Montipora* (Figure 6I, *rectangle dashed lines*) (Supplementary Movie 9)

Together, our observations provide direct evidence that coral colonies can form integrated systems, in which some individual polyps are physically connected through a continuous gastrovascular network. This internal connectivity likely facilitates the exchange of nutrients, metabolites, and signaling molecules, linking polyp-level organization to colony-level function.

### Coral colony morphology and evolution

Having characterized polyp morphology and internal connectivity, we next sought to determine how these soft tissue traits relate to the skeletal structures that house them. To do this, we mapped the position and organization of polyps onto their corresponding corallites across the different coral species. Using *A. cervicornis* as a reference, we defined key components of coral skeletal architecture, including the theca, corallite, and coenosarc (Figure 7A). Comparative surface views revealed species-specific differences in polyp arrangement and colony morphology, ranging from the structured axial–radial organization in *A. cervicornis* to the more uniform *A. millepora*, the dense packing of *P. damicornis*, and the embedded polyp configuration of *M. capitata* (Figure 7B–D).

**Figure 7.**
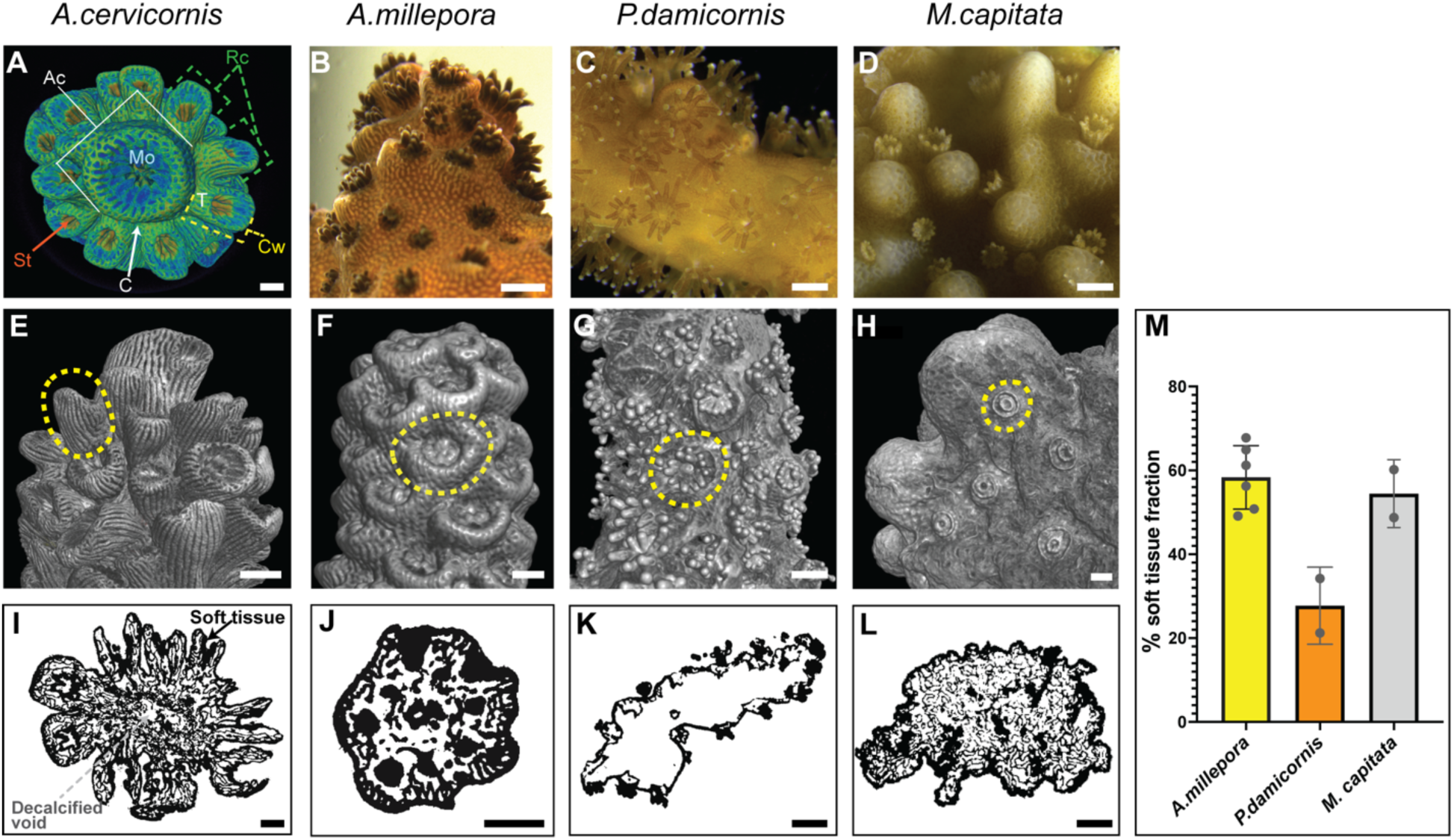
External morphology and corallite architecture across four coral species. **A**, Top view of *A. cervicornis* branch tip highlighting the axial corallite (*A*c) surrounded by radial corallites (*Rc*). Structural features of the axial corallite are indicated, including the mouth (*Mo*), tentacles (*T*), septa (*St*), coenosarc (*C*), and corallite wall (*Cw*). **B–D,** Live colony views showing polyp organization at branch tips. **B,** *A. millepora* branch tip showing the arrangement of polyps along the branch surface. **C,** *P. damicornis* colony surface with multiple extended polyps and visible tentacles. **D,** *M. capitata* colony surface showing small polyps embedded within the coenosteum between skeletal mounds. **E–H,** Volume rendering of superficial structures corresponding to the species shown above, illustrating corallite morphology and skeletal arrangement. Dashed circles highlight representative corallites for each species. (E) *A. cervicornis* showing the axial corallite and surrounding radial corallites. (F) *A. millepora* with a prominent corallite structure along the branch surface. (G) *P. damicornis* showing densely packed corallites distributed across the colony surface. (H) *M. capitata* showing embedded corallites within the coenosteum. **I–L**, Representative binary cross sections used for tissue fraction analysis. Black regions indicate soft tissue, and white regions indicate space left after the hard skeleton was removed by decalcification. Panels show (I) *A. millepora*, (J) *A. cervicornis*, (K) *P. damicornis,* and (L) *M. capitata*. (M) Column plot showing soft tissue volume fraction among species. Data are presented as means. Mean soft tissue volume fraction was 58.35 % (*n*= 6) in *A. millepora*; 54.46 % (*n*= 2) in *M. capitata*; 27.72 % (*n*= 2) in *P. damicornis*. Statistical analysis was performed using the Wilcoxon test in GraphPad Prism. Scale bars, 1000μm.

Volume rendering further highlighted differences in corallite architecture across species (Figure 7E–H). In *A. cervicornis* and *A. millepora*, possessed prominent tubular corallites projecting from the colony surface. In contrast, *M. capitata* and *P. damicornis* colonies exhibited low-relief, ring-like corallites embedded within the coenosteum. To characterize species-specific differences in internal organization, we analyzed the soft tissue volume fraction, which represents how much of the analyzed fragment is occupied by soft tissue. In this analysis, black regions corresponded to soft tissue while white regions represented the void left after decalcification (Figure 7I–L). *A. cervicornis* exhibited a relatively continuous tissue organization interspersed with irregular lumen spaces (Figure 7I). *A. millepora* showed a compact and cohesive tissue matrix with smaller, interconnected lumen spaces (Figure 7J). In contrast, *P. damicornis* revealed large, open lumen regions with reduced tissue connectivity (Figure 7K), whereas *M. capitata* showed a densely packed tissue organization with a fine network of small lumen spaces (Figure 7L).

We next performed quantification of binary cross sections and found differences in soft tissue volume fraction among species (Figure 7M; Supplemental File 4). Soft tissue volume fraction measurements were derived from a limited number of fragments per species. *A. millepora* exhibited the highest mean soft tissue fraction (58.35 %, *n* = 6), followed by *M. capitata* (54.46%, *n* = 2), whereas *P. damicornis* showed a markedly lower value (27.72%, *n* = 2). Thus, *P. damicornis* contained proportionally less soft tissue, leaving a larger fraction as decalcified skeletal space. Hence these measurements are best interpreted as estimates of internal architectural diversity. Larger comparative datasets would be needed to establish statistically validated differences across species. Nevertheless, these measurements provide an approximation of the relative proportion of tissue to internal void space, highlighting a range from lumen-dominated to tissue-dense organizations. Collectively, these findings demonstrate that variation in internal tissue architecture and skeletal housing underlies differences in coral growth form, functional integration, and ecological strategy.

From an evolutionary perspective, our observations raise the question of whether variations in internal polyp organization reflect independent adaptations to distinct ecological conditions. The marked divergence of internal structure between *Acropora* and *Montipora*, despite their shared family membership within Acroporidae, suggests that internal polyp architecture may not strictly follow phylogenetic relatedness. To explore it, we mapped our morphological findings onto a time-calibrated phylogeny reconstructed from mitochondrial cytochrome *b* (CytB) nucleotide sequences (Figure 8). The resulting tree was consistent with established relationships among major cnidarian lineages and confirmed the monophyly of Scleractinia, with an estimated origin around 240–250 million years ago. Although *A. cervicornis*, *A. millepora*, and *M. capitata* are more closely related within Acroporidae, they exhibit substantial differences in polyp hierarchy, tissue organization, and gastrovascular connectivity. Conversely, some structural features are shared across more distantly related taxa, indicating that aspects of internal organization may evolve independently of phylogenetic proximity.

**Figure 8.**
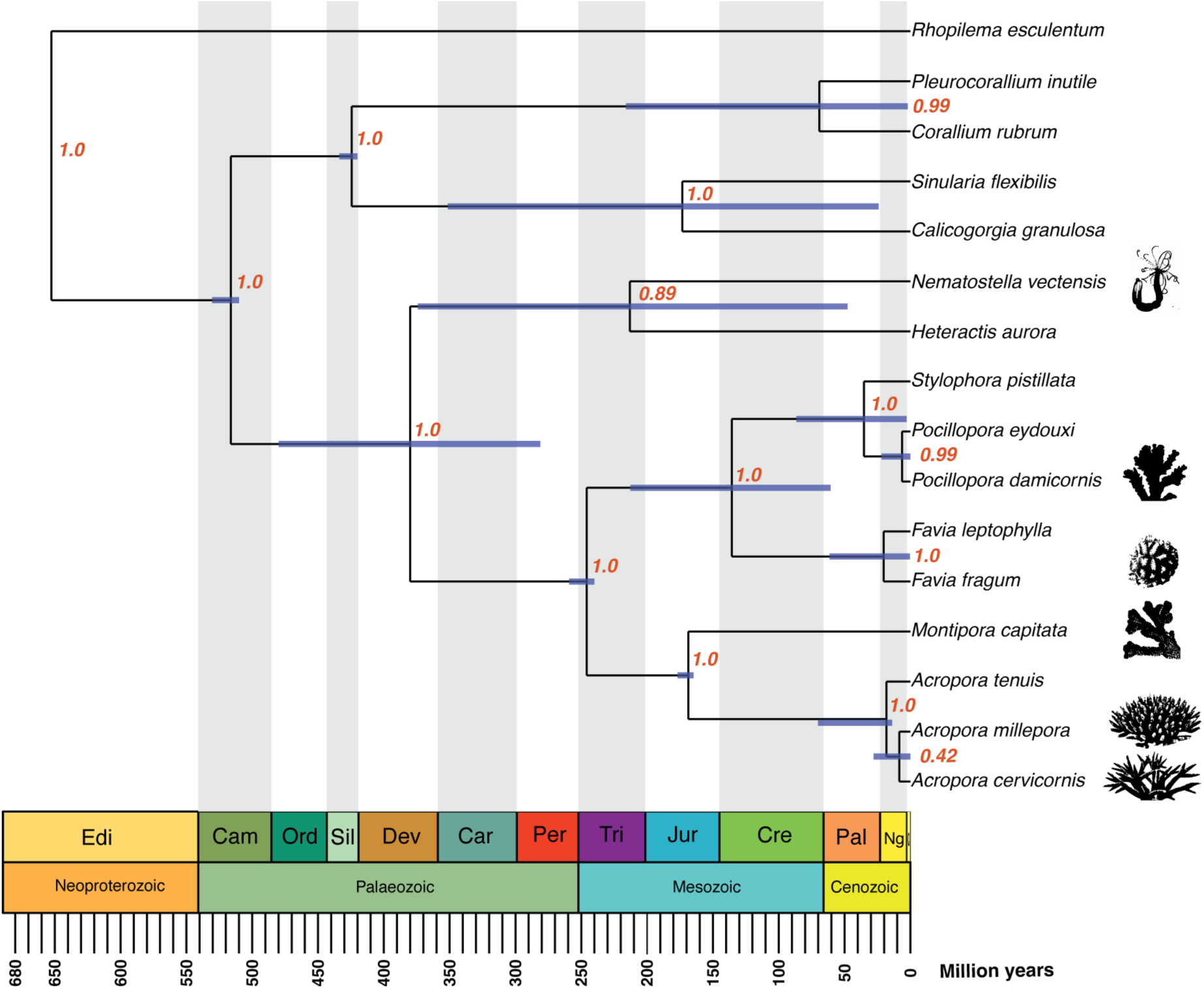
Time-calibrated phylogenetic framework of selected cnidarian taxa based on cytochrome *b* (CytB) nucleotide sequences. The tree places the focal scleractinian coral species analyzed in this study (*Acropora cervicornis, Acropora millepora, Montipora capitata,* and *Pocillopora damicornis*) within a broader cnidarian evolutionary context. Major clades, including *Medusozoa* and *Anthozoa*, were resolved, with Scleractinia recovered as a monophyletic group. Within Scleractinia, acroporid taxa (*Acropora spp*. and *Montipora capitata*) cluster together, whereas *Pocillopora* and related taxa form a distinct lineage. Branch lengths are proportional to time, with divergence estimates mapped onto the geological timescale from the Ediacaran to the present (bottom axis). Blue horizontal bars at nodes indicate 95% highest posterior density (HPD) intervals for divergence times, and red values represent posterior probabilities supporting each node. The position of the focal species provides an evolutionary framework for interpreting variation in internal polyp organization and tissue architecture across taxa.

Together, our results suggest that coral diversity encompasses not only variation in skeletal morphology but also significant divergence in internal tissue architecture. The mismatch between phylogeny and internal organization suggests that internal tissue architecture is evolutionarily plastic, shaped as much by developmental and ecological pressures as by shared ancestry.

## Discussion

### Resolving coral soft tissue architecture in 3D

Our 3D reconstructions demonstrate that coral diversity extends beyond external colony morphology, uncovering a previously difficult-to-access layer of internal organization defined by polyp architecture, tissue distribution, and gastrovascular connectivity in stony corals. By combining contrast-enhanced X-ray tomography (XRT) imaging with three-dimensional reconstruction, we resolved features of internal colony structure not captured by traditional X-ray or surface-based analyses.

Comparing internal organization across coral species (including polyp hierarchy, mesenterial length and complexity, tissue distribution, and gastrovascular connectivity) revealed substantial variation that did not map neatly onto phylogenetic relationships. For example, the three acroporid species in our study each displayed distinct internal arrangements despite belonging to the same family, while certain traits such as reduced mesenterial length appeared in both *P. damicornis* and *A. millepora* regardless of their evolutionary distance (Figure 5C). As with any contrast-enhanced XRT approach, differences in decalcification duration, tissue response to fixation, and contrast agent penetration across species could influence the appearance of internal structures to some degree. Further standardization of preparation protocols would help refine these comparisons. Nevertheless, the consistency of observed patterns across multiple independent traits and species, supports their interpretation as genuine biological variation. Furthermore, this data suggests that internal coral tissue architecture represents an evolutionarily flexible component of coral biology that is not fully captured by traditional skeletal-based classifications.

Traditionally, X-ray tomography and its derivatives have focused on skeletal morphometrics and microarchitecture ^7, 9–11, 13–16, 34^, while the organization of living tissues in stony corals has remained largely unexplored due to their dense mineralized skeleton. Our workflow addresses this gap, but several limitations should be acknowledged. Sample availability was constrained for some species, which prevented robust statistical comparisons of metrics such as void volume fraction. Tissue fixation caused contraction of soft structures, in particular tentacles, likely leading to underestimation of polyp dimensions. And while threshold-based segmentation enabled visualization of tissue and lumen compartments, it may introduce uncertainty when distinguishing fine-scale structures. Expanding sample sizes across species and functional complementary approaches such as live imaging, fluid flow modeling, or tracer-based experiments, will be essential to functionally link the internal architectures described here with physiological function.

Despite their central role as reef builders, resolving the internal organization of scleractinian corals remains challenging. Their dense aragonitic exoskeleton ^18, 19^, imposes a major limitation for X-ray–based imaging, as its strong absorption reduces soft tissue contrast in XRT, thereby constraining the resolution of internal anatomical features. Furthermore, their thin and fragile soft tissue is prone to collapse and morphological loss during decalcification. As a result, traditional two-dimensional histology remains the standard for examining internal anatomy ^20, 21^ but fails to resolve three-dimensional connectivity within the colony.

In order to resolve the internal organization of corals, several approaches have focused on soft corals (*Octocorallia*), which also contribute to reef formation and often exhibit greater resilience to environmental stressors ^35^. These organisms possess a dynamic skeletal system of mineralized sclerites embedded within a mesoglea gel matrix, providing mechanical properties, including stiffness and strength ^36^. Their less dense and polymorphic framework of aragonite and calcite ^37^, enable direct contrast-enhanced imaging of soft tissues without decalcification ^22, 38^.

X-ray Tomography (XRT) has been broadly used to visualize mineralized tissues and has recently adapted for soft-tissue imaging through contrast-enhanced approaches. Lugol’s iodine and phosphotungstic acid (PTA)–based solutions are among the most the most commonly used contrast agents due to their relative accessibility and lower toxicity, generating distinct patterns of X-ray attenuation ^29, 30^. Iodine-based solutions provide broad contrast enhancement and have been successfully applied in octocorals ^22^, whereas PTA preferentially stains protein-rich connective tissues such as muscles and internal structures ^29, 31^.

Although decalcification combined with contrast-enhanced XRT imaging is well established for imaging mineralized tissues in other biological systems ^31, 39, 40^, its application in scleractinian corals remains limited by restricted penetration and interference from the mineralized skeleton. Here, we optimized a contrast-enhance workflow to visualize the internal tissue architecture, where the dense aragonite skeleton has long obscured internal tissue organization from conventional imaging.

### Internal architecture and the evolution of coral form: a framework linking development, ecology, and diversification

The evolution of modular colonial organisms such as reef-building corals is intriguing and complex due to their dual reproductive strategy, in which modules (e.g., polyps or polyps together with their skeletal corallite) proliferate asexually while whole colonies reproduce sexually. This duality allows selection to operate across hierarchical levels, potentially generating conflicting evolutionary pressures between modules and colonies ^3, 4^. Evidence from bryozoans such as *Stylopoma* demonstrates that module-level traits exhibit variability within colonies but are not strongly heritable across generations. In contrast, colony-level traits, defined by the spatial organization, integration, and coordination of modules, are consistently inherited and represent the primary targets of natural selection ^4^. Because individual modules do not transmit their traits to subsequent modules, selection at the modular level may be constrained, whereas colony-level traits are more evolvable.

Colonial organisms function as highly integrated biological systems. In reef-building corals, this integration is reflected variation of colony-level traits such as branching architecture and spatial organization, which vary with environmental conditions ^5^. This structural complexity arises from interplay between intrinsic genetic programs and environmental plasticity, reflecting a dynamic balance that governs coral colony morphogenesis ^5^.

A well-characterized example of this hierarchical integration is provided by *Acropora* species, in which polyps are interconnected through a highly structured, mesh-like canal network and conserved axial-radial pattern ^34^. In this context, the axial axis plays a central role as both a directional organizer of colony growth and a conduit for internal nutrients. This organization is associated with internal hydrodynamics, where axial flow dominates in the upper regions, supporting rapid apical extension, while radial flow in lower regions contributes to skeletal thickening and mechanical stability ^8, 41^. These observations suggest that the axial-radial architecture of *Acropora* integrates growth, transport, and structural reinforcement within a single colony framework.

Our observations directly support this hierarchical organization at the level of polyp morphology; we identified species-specific variation in axial polyp morphology. In *A. cervicornis*, the axial polyp displayed asymmetric, elongated mesenteries extending deeply into the axial lumen, whereas in *A. millepora*, the axial polyp exhibited a more homogeneous mesenterial length that did not extend as deeply along the main axis; these differences, potentially reflect distinct patterns of internal transport and growth regulation.

Importantly, these colony-level morphologies emerge from local interactions among polyps ^5^, emphasizing the need to incorporate polyp-level traits into an eco–evo–devo perspective. Despite not directly driving evolution, polyp-level traits define the developmental rules and constraints that generate heritable colony-level architectures. In this context, polyp morphology represents a critical interface between development, physiology, and evolution ^5, 42^.

### Species-specific polyp connectivity strategies in corals reflect architecture-dependent modes of internal communication and integration

Coral colonies have an integrated nutrient distribution system, connecting the colony surface, neighboring polyps and internal tissues, where surface and internal flows jointly regulate resource distribution and growth ^26, 27^. Ciliated cells on the coral surface enhance exchange at this interface, whereas tissue continuity and gastrovascular connectivity may facilitate the redistribution of nutrients across the colony and toward more internal regions ^24, 25^. These findings indicate that ciliary activity is a critical component of coral physiology, linking polyp-level processes to colony-scale function and potentially contributing to the ecological success of reef-building corals ^25, 28^.

A useful comparison comes from hydroids, which are colonial cnidarians that connect their polyps through specialized tubular structures called stolons. These root-like connections anchor the colony to the substrate while serving as conduits for internal fluid transport, supporting digestion, nutrient distribution, and asexual reproduction ^23, 27, 43^. What makes this system particularly interesting is that at polyp–stolon junctions, mitochondrion-rich myoepithelial cells act as valves, actively regulating gastrovascular flow through the colony ^23^. Although corals lack stolons, the mesenterial connections identified here may serve an analogous role, physically bridging gastrovascular cavities and allowing fluids and nutrients to move between polyps. The direct axial-radial connections in *Acropora* and the multi-polyp networks in *M. capitata* together suggest that scleractinian corals have evolved distinct structural solutions for integrating the colony, despite fundamental anatomical differences from hydroid stolon systems. Whether coral mesenterial junctions contain specialized cell types like the myoepithelial cells in hydroids remains an open question that will require ultrastructural work to resolve.

Notably, species-specific differences in morphology appear to dictate modes of interaction and connectivity. As mentioned above, in *Acropora*, the dominance of the axial axis highlights a direct linkage between the axial polyp and surrounding radial polyps, potentially mediated through mesenterial extensions that traverse the axial lumen. In newly forming polyps of the same individual, connectivity may occur through superficial coenosarc tissue, providing an initial integrative framework prior to the establishment of fully developed gastrovascular connections. In *Montipora capitata*, polyp connectivity does not appear to reflect a simple proximity-based spatial pattern; instead, connectivity appears heterogeneous, with individual polyps forming multiple links with both adjacent and more distant neighbors. In contrast, *P. damicornis* displayed superficial connectivity just below the coenosarc, rather than intricate network extending through the main lumen. This observation is consistent with previous studies indicating that living polyps are restricted to the colony surface. Likewise, 3D skeletal reconstruction showed that each polyp occupied a calice with no direct connection to adjacent calices ^11^. However, our observations suggest that, despite this skeletal separation, polyps were connected through mesenterial association. This variability suggests that canal connectivity is not solely determined by spatial arrangement but may instead be governed by intrinsic developmental or physiological rules. Interestingly, work on *Pocillopora grandis*, has recently demonstrated that corallite microstructure changes with developmental stage and season and is not the fixed trait it was long assumed to be ^44^. This means the internal organization we observed in *P. damicornis* could partly reflect the developmental stage at which the colony was collected.

Colony morphology arises from the interaction between physiological demands and environmental constraints, optimizing resource acquisition and coral architecture under different ecological conditions. Although *Acropora* and *Montipora* are both primary reef-builders within the family Acroporidae, they occupy distinct ecological niches and exhibited striking differences in colony architecture, polyp morphology, internal tissue organization. From an eco–evo–devo perspective, this contrast illustrates that closely related corals can evolve divergent developmental and architectural strategies in response to their environments. This interpretation is further supported by the phylogenetic analysis, which showed that internal organization does not closely track evolutionary relatedness. Despite belonging to the same family, *Acropora* and *Montipora* displayed markedly different connectivity patterns, whereas the distantly related *A. millepora* and *P. damicornis* converged on more compact polyp architectures. This suggests that internal organization represents an evolutionarily flexible trait, shaped by developmental and ecological processes rather than strictly constrained by phylogenetic history ^4, 5^.

Finally, our findings raise fundamental questions regarding the mechanisms that regulate polyp–polyp communication, including whether specialized mesenterial structures or alternative pathways mediate these connections. In this sense, structural connectivity might represent a key unexplored feature underlying colony integration in reef-building corals, one that might contribute to a better understanding their development, physiology and environmental change.

## Materials and Methods

### Sample collection

Coral fragments were obtained from a closed-system mesocosm maintained under controlled environmental conditions that simulated natural reef temperature, photoperiod, solar irradiance, and lunar cycles ^45^. The only exception was *Favia gravida*, which was acquired from a local aquarium supplier. Prior to fixation, all specimens were maintained in filtered seawater at 35 ppt salinity.

Fragment dimensions were estimated before fixation and are reported as length × width × height. The approximate dimensions were as follows: *A. cervicornis*, 3.8 x 2.3 × 1.2 cm; *M. capitata*_*1*, 3.0 × 2.3 × 1.5 cm; *M. capitata_2*, 2.7 × 1.1 × 0.9 cm; *P. damicornis_1*, 3.8 × 2.0 × 0.8 cm; *P. damicornis_2*, 3.1 × 1.3 × 0.7 cm; *A. millepora_1.1–1.3*, 5.0 × 2.5 × 1.2 cm; *A. millepora_2.1–2.3*, 5.4 × 2.6 × 1.3 cm; *A. millepora_3_1*, 1.6 × 0.8 × 1.1 cm; *A. millepora_3_2*, 3.0 × 0.6 × 1.2 cm; *A. millepora 3_3*, 4.1 × 0.7 × 1.2 cm; and *Favia gravida*, 3.0 × 1.8 × 2.0 cm.

Samples were anesthetized with approx. 100-200 μl of 7% MgCl_2_ in 35 ppt seawater for 15 min to relax the coral tissues and promote tentacle extension. However, despite MgCl_2_ anesthesia, tentacle contraction was observed after the addition of the 4% PFA fixative solution. Most of the seawater was then removed, and specimens were fixed in 4% paraformaldehyde diluted in seawater at 4°C overnight on a shaker. Following fixation, samples were washed five times in 1x PBS and subsequently decalcified with EDTA.

### Conventional X-ray Tomography (XRT)

This protocol was adapted from established methods for visualizing soft tissue within mineralized tissues of mouse ^46, 47^ and human ^48^ samples, as well as previously described procedures for soft tissue XRT ^31, 49^. Briefly, samples were decalcified with 0.5M EDTA (see Supplementary Table 1 for detailed conditions), (EDTA solution was replaced every three days). Following decalcification, samples were rinsed three times with sterilized PBS (pH 7.2) for 10 min on a slow-speed orbital shaker. Dehydration was performed by a graded ethanol series: 30%, 50%, and 70% (v/v) for 1 h per step. Samples were then immersed in 1% (w/v) Phosphotungstic Acid (Electron Microscopy Science, 19500) in 70% ethanol for contrast enhancement. The staining solution was refreshed weekly, with a total staining duration of around four weeks.

Following staining, samples were briefly rinsed in 70% ethanol and immediately immobilized in 1% low melting-point agarose (maintained at 42°C in a water bath) within a 1 mL or 5 mL microfuge tube or a custom 3D-printed plastic tube depending on the size of the sample. Once the agarose solidified, the specimen was then mounted onto a custom designed 3D-printed scanning sample holder using paraffin and parafilm. 3D print files for custom tubes and holders can be found in Supplementary Files S1, S2. Conventional XRT imaging was performed using a SkyScan 1272 system (Bruker), with the specific scanning parameters summarized in Supplementary Table 1. The resulting 2D projections were reconstructed using NRecon software (v2.2.0.6, Bruker), and 3D image analysis was conducted using Amira software (v2024.2, Thermo Fisher Scientific). Dragonfly software (v2025.1; Object Research Systems, ORS Inc., Montreal, Canada) was additionally used for image visualization, segmentation, 3D reconstruction, and rendering of three-dimensional image sequences.

### X-ray microscopy (XRM)

A subset of samples prepared for conventional XRT were also imaged using X-ray microscopy (XRM), using a ZEISS Xradia 520 Versa. Contrast enhanced samples in agarose were mounted on XRM sample holders for multiple scans using a range of objective lenses, allowing multiscale imaging to place high magnification scan data within the context of the entire sample. Scan parameters are shown in Supplementary Table 2. Scan data were reconstructed into 3D volume data using ZEISS Reconstructor software, and exported as 16-bit TIFF stacks for analysis as described.

### Statistical analysis

Statistical analyses were performed using GraphPad Prism software. Differences among groups were evaluated by one-way analysis of variance (ANOVA). Data are presented as mean ± SD, and statistical significance was set at P < 0.0001. The measurements used to calculate total length and individually segmented polyps are provided in Supplementary Table 3.

### Volume fraction analysis

Volume fraction analysis was used to quantify the relative proportions of soft tissue and internal void (lumen) space within the reconstructed volumes. Decalcified corals were segmented in 3D using Segment Anything 2 ^50^. A single frame was annotated by drawing a box, the coral identified and that segmentation propagated throughout the image stack. To segment the soft tissue: XRT and XRM data were converted to absorbance, background subtracted, gaussian blurred and thresholded with a constant absolute absorbance across all samples that was chosen to avoid noise and include obvious tissue. This thresholded image was masked by the whole coral segmentation, and the total volumes of each segmentation calculated. The soft tissue volume was divided by the total coral volume to obtain the soft tissue volume fraction for each samples. This analysis was done in python using scipy, scikit-image ^51^ and napari ^52^ (Supplementary Table 4).

### Sequences, alignment, phylogenetic inference and molecular dating

Nucleotide sequences of the mitochondrial cytochrome *b* (*cytb*) gene were obtained from publicly available databases (e.g., GenBank, NCBI) for representative taxa of Scleractinia, Octocorallia, Actiniaria, and Medusozoa. Taxon sampling was designed to capture major cnidarian lineages and include both focal and outgroup taxa. The sequences are from: *Rhopilema esculentum* (GenBank KY454768.1), *Pleurocorallium inutile* (GenBank LC848439.1), *Corallium rubrum* (GenBank AB700136.1), *Sinularia flexibilis* (NCBI NC_061282.1), *Calicogorgia granulosa* (NCBI NC_023345.1), *Nematostella vectensis* (GenBank OW052000.1), *Heteractis aurora* (NCBI NC_047219.1), *Stylophora pistillata* (NCBI NC_011162.1), *Pocillopora eydouxi* (NCBI NC_009798.1), *Pocillopora damicornis* (GenBank ACA84081.1), *Favia leptophylla* (also known as *Mussismilia leptophyllai* NCBI AB117307), *Montipora capitata* (GenBank MK091602.1), *Acropora tenuis* (NCBI LC201868), *Acropora millepora* (GenBank AF099653.1), *Acropora cervicornis* (GenBank MW246492.1). Sequences were aligned using standard multiple sequence alignment approaches (MAFFT) and alignments were manually inspected and edited to ensure positional homology and remove ambiguous regions with SEAVIEW. Only high-confidence aligned regions were retained for downstream analyses.

Bayesian phylogenetic inference and divergence time estimation were conducted using BEAST 2 ^53^. The best-fitting nucleotide substitution model was selected prior to analysis using established model-selection frameworks by jModelTest ^54^. Among-site rate heterogeneity was modeled using a gamma distribution, then a relaxed uncorrelated lognormal molecular clock was implemented to accommodate substitution rate variation among lineages ^55^. The branching process was modeled under a Yule speciation prior, which is appropriate for interspecific phylogenetic datasets (Supplementary Table 5 & Supplementary Data 1).

### Fossil calibrations

Divergence time estimation was calibrated using fossil-based minimum age constraints applied as lognormal priors on selected nodes. Calibration points were chosen based on fossil evidence and previous molecular clock studies. The crown group of Scleractinia was constrained with a minimum age of 240 Ma (Triassic) ^56^. The crown group of Acroporidae was constrained with a minimum age of 150–160 Ma (Jurassic), consistent with fossil representatives of acroporid-like ^57^. The crown group of Octocorallia was constrained with a minimum age of 420 Ma (Silurian), based on early octocoral-like fossils and molecular estimates ^58^.

The clade Medusozoa was constrained with a minimum age of 510 Ma (Cambrian), reflecting the earliest fossil evidence of jellyfish-like organisms ^59^. All calibrations were implemented as lognormal priors, with offsets corresponding to minimum fossil ages and standard deviations (S) and mean parameters (M) chosen to allow soft upper bounds, thereby accommodating uncertainty in the fossil record while enforcing minimum constraints. Markov chain Monte Carlo (MCMC) analyses were run for 50 million generations, sampling every 5,000 generations. Convergence and mixing were assessed using Tracer ^60^ where all ESS parameters exceeded 200. A burn-in of 10% of the sampled trees was applied to exclude the initial non-stationary phase of the MCMC (Supplemental File S5).

## Supporting information

Supplementary File S1_holder

Supplementary File S2_tube

Supplementary Movie 1_A.cervicornis

Supplementary Movie 2_A.millepora

Supplementary Movie 3_P.damicornis

Supplementary Movie 4_M.capitata

Supplementary Movie 5_A.cervicornis top view

Supplementary Movie 6_M.capitata

Supplementary Movie 7_A.millepora link

Supplementary Movie 8_M.capitata gastrovascular connectivy

Supplementary Movie 9_P.damicornis gastrovascular connectivy

## Acknowledgements

We thank the Invertebrate Research Organisms Technology Center at Stowers Institute for Medical Research, especially Shane Miller, Rachel Livella and Jason Livella, and the Invertebrate team for their support obtaining coral samples and maintaining the organisms.

This study was funded through the generous support of the Stowers Institute for Medical Research.

## Authorship contributions

**Emma Rangel-Huerta**: Conceptualization, Investigation, Methodology, Project administration, Visualization, Writing – original draft, Writing – review & editing, Validation. **Meiru Wang**: Conceptualization, Investigation, Methodology, Visualization, Resources, Writing – review & editing. **Stephanie H. Nowotarski**: Investigation, Methodology, Visualization, Resources, Writing – review & editing. **Keith Duncan**: Visualization, Resources, Writing – review & editing. **Sean McKinney**: Data curation, Formal analysis, Methodology, Software, Writing – review & editing. **Matthew C. Gibson**: Funding acquisition, Project administration, Supervision, Writing – review & editing. All authors have read and agreed to be listed on this manuscript.

## Competing interests

The authors have no conflict of interests to disclose

## Declaration of generative AI and AI-assisted technologies in the writing process

During the preparation of this work, the author(s) used the assistant of ChatGPT to assist with grammar and spelling. After using this tool, the author(s) review and edited the content as needed and take full responsibility for the final content of the manuscript.

## Notes

### Competing Interest Statement

The authors have declared no competing interest.

